# Dysregulated expanded endocannabinoid system as therapeutic targets of amyotrophic lateral sclerosis

**DOI:** 10.1101/2024.01.12.575341

**Authors:** Daisuke Ito, Madoka Iida, Yohei Iguchi, Atsushi Hashizume, Shinichiro Yamada, Yoshiyuki Kishimoto, Shota Komori, Teppei Shimamura, Yuto Takemoto, Masahiro Nakatochi, Tomohiro Akashi, Kunihiko Hinohara, Hyeon-Cheol Lee-Okada, Junichi Niwa, Gen Sobue, Shinji Tanaka, Ken Takashina, Takehiko Yokomizo, Masahisa Katsuno

## Abstract

Amyotrophic lateral sclerosis (ALS) is a devastating neurodegenerative disease characterized by the selective loss of upper and lower motor neurons. ALS patients often manifest systemic metabolic abnormalities such as glucose intolerance. Herein, to elucidate the systemic metabolic changes related to ALS progression, we performed metabolomics analysis on the serum of ALS patients and identified several metabolites associated with the disease progression, including metabolites involved in the expanded endocannabinoid system (ECS). In particular, the levels of N-acyl taurines (NAT) were correlated with the longitudinal change in the revised ALS functional rating scale (ALSFRS-R) rating. In vitro experiments with ALS cell models and in vivo studies with SOD1^G93A^ transgenic mice revealed that PF-04457845, a fatty amide acid hydrolase (FAAH) inhibitor, up-regulated the expanded ECS, particularly the levels of NATs and N-acyl ethanolamine and ameliorates motor neuron degeneration through the regulation of microglial polarization, synapse plasticity, and neuronal development. Our study indicates that dysregulation of the expanded ECS is associated with ALS progression and a target for novel disease-modifying therapies.

## INTRODUCTION

Amyotrophic lateral sclerosis (ALS) is a devastating neurodegenerative disease caused by the selective loss of upper and lower motor neurons. Approximately 95% of ALS cases are sporadic, and 5% are familial. Causative genes for familial ALS are diverse, including *superoxide dismutase 1* (*SOD1*), *TARDBP*, and *C9orf72* (Yokoseki *et al*, 2008; Sreedharan *et al*, 2008; Kabashi *et al*, 2008; Gitcho *et al*, 2008; Renton *et al*, 2011; DeJesus-Hernandez *et al*, 2011; Rosen *et al*, 1993). A *SOD1*-based murine model has been widely used for the preclinical study of ALS therapeutic strategies because it simulates the rapid progression of motor neuron degeneration and glial cell changes characteristic of human ALS pathology. In sporadic ALS, muscle weakness and atrophy rapidly progress, and the time from onset to death is around 3–5 years, mainly as a result of respiratory failure (Wijesekera & Leigh, 2009; Rowland & Shneider, 2001). However, it is widely known that there is a considerable variability in the disease progression of sporadic ALS. This variability suggests that biological factors governing disease progression may be a therapeutic target of sporadic ALS. However, the molecular basis for ALS phenotypic heterogeneity remains elusive.

A great number of therapeutics for ALS that showed efficacy in preclinical studies failed in clinical trials (Mitsumoto *et al*, 2014; Petrov *et al*, 2017). Although riluzole and edaravone have been approved for ALS treatment, their efficacies are limited(Bensimon *et al*, 1994; Abe *et al*, 2017). These difficulties in therapy development are, at least partially, due to the discrepancy between the molecular changes in human sporadic ALS and those in cell or animal models established by implementing familial ALS gene mutations; the cell and animal models expressing the causative gene mutations of familial ALS do not necessarily reflect the pathological mechanism of human sporadic ALS. Therefore, the development of therapeutics should be based on human ALS pathophysiology. An approach to overcome these struggles is the utilization of induced pluripotent stem (iPS) cells derived from human ALS patients (Baxi *et al*, 2022). Indeed, drug screening using motor neurons differentiated from iPS cells identified several candidates for ALS therapeutics (Imamura *et al*, 2017; Fujimori *et al*, 2018). However, with this approach, it is difficult to target glial pathology, which plays fundamental roles in ALS progression (Van Harten *et al*, 2021; Boillée *et al*, 2006; Yamanaka *et al*, 2008). Thus, more comprehensive approaches are needed to achieve reverse translational therapy development based on human ALS pathophysiology.

Patients with sporadic ALS often develop a pathological condition called hypermetabolism, a severe weight loss due to excess resting energy expenditure, which is related to rapid progression and poor prognosis of ALS (Bouteloup *et al*, 2009; Steyn *et al*, 2018; Jésus *et al*, 2018). In addition, glucose intolerance and dyslipidemia have also been reported in ALS patients, even at a prodromal stage of disease (Mariosa *et al*, 2017; Araki *et al*, 2019). Nutritional intervention with a high-calorie and/or high-fat diet suppresses disease progression in certain populations of sporadic ALS patients (Ludolph *et al*, 2020; Wills *et al*, 2014). These observations suggest a causative role of systemic metabolic changes in ALS.

Herein, we aimed to develop a therapeutic strategy for ALS with a special focus on metabolic alterations in patients by reverse translating clinical observations to basic research. We first conducted a clinical study to elucidate metabolic changes related to disease progression, screened compounds targeting the identified metabolic profiles in cellular models, and then tested hit compounds in a mouse model of ALS. Our study identified that the dysregulation of the expanded endocannabinoid systems (ECS) such as *N*-acyl taurines (NATs) is associated with disease progression in patients with sporadic ALS and that a pharmacological approach that activates the expanded ECS is a potential therapeutic strategy for ALS.

## RESULTS

### Serum metabolome analysis in sporadic ALS patients

The serum metabolome was analyzed in subjects with sporadic ALS and healthy controls. A total of 26 subjects with ALS and 10 healthy controls were analyzed (Fig. 1A). The median longitudinal change in the revised ALS Functional Rating Scale (ALSFRS-R) at 6 months (ΔALSFRS-R) was –4. Therefore, subjects with ALS whose ΔALSFRS-R for 6 months was less than –4 were defined as having rapidly progressive ALS (Rapid ALS, n = 12), and the others were defined as having slowly progressive ALS (Slow ALS, n = 14). Baseline characteristics were not different between the subjects in the Rapid ALS and Slow ALS groups (table S1). Sera of ALS patients and healthy controls were subjected to metabolome analysis with HPLC MS/MS.

**Fig. 1.**
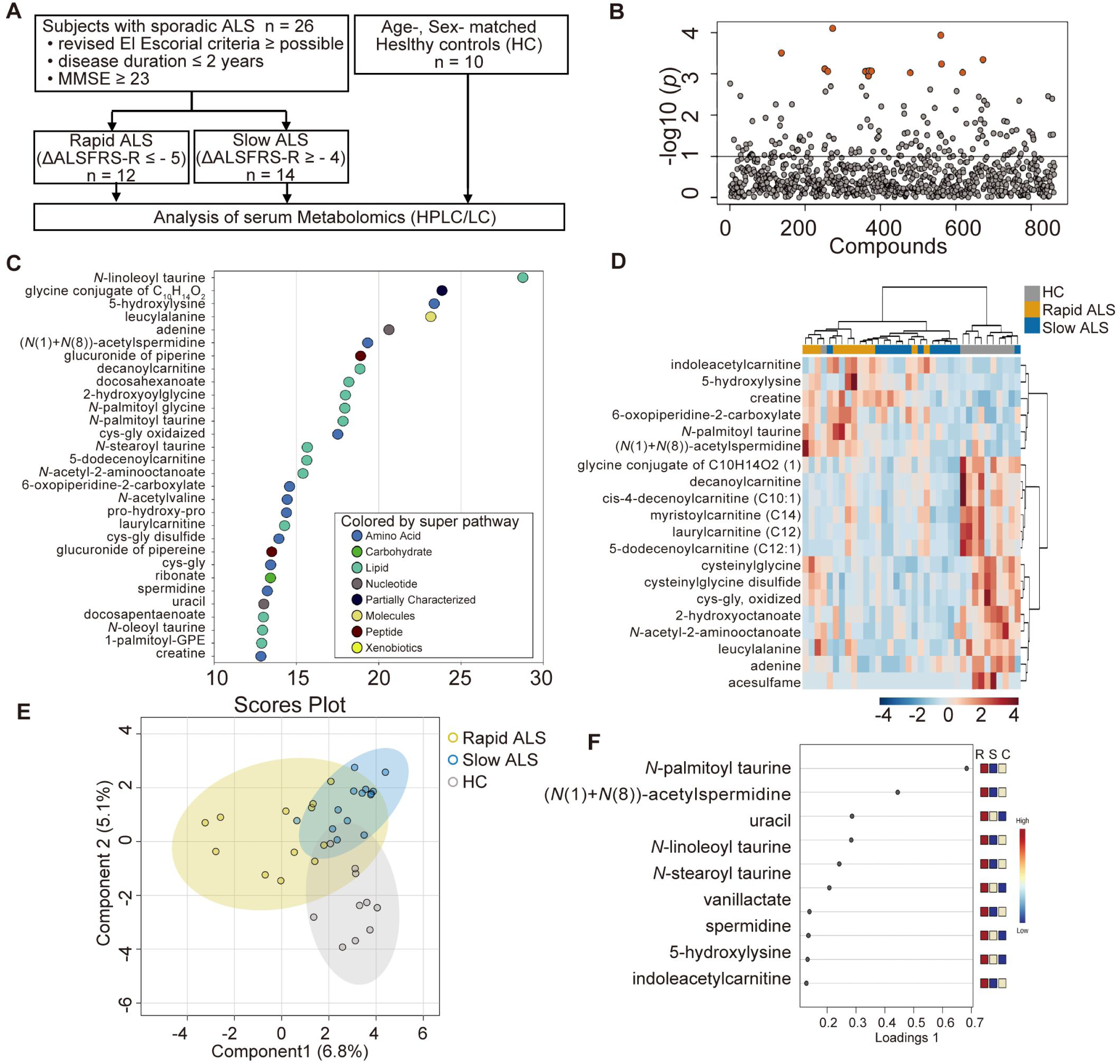
Metabolomics analysis of patients with ALS and healthy individuals. (**A**) Flow chart of metabolome analysis. We included subjects with definite, probable, or possible ALS that met the revised El Escorial criteria and had disease duration less than 2 years and an MMSE score of 23 or higher. Subjects with ALS were divided into rapidly progressive (Rapid ALS) and slowly progressive (Slow ALS) groups according to the median longitudinal change in ALSFRS-R from the first evaluation to 6 months. We also included 10 age– and sex-matched healthy controls. (**B**) Manhattan plot of metabolomics by UPLC‒MS/MS. Serum metabolites with different levels among Rapid ALS, Slow ALS, and healthy controls determined by MetaboAnalyst 5.0 are shown in orange circles (FDR > 0.1, one-way ANOVA). (**C**) Top 30 metabolites that discriminate among Rapid ALS, Slow ALS, and healthy controls by random forest classification. The metabolites were ranked according to their mean decrease accuracy. (**D**) Heatmap analysis of the top 20 metabolites identified by one-way ANOVA. (**E**) Sparse partial least squares discriminant analysis (sPLS-DA) plots by MetaboAnalyst 5.0 to discriminate Rapid ALS, Slow ALS, and healthy controls. (**F**) Loading plots of the top 10 metabolites by sPLS-DA shown in (E).

Serum metabolomics detected 867 metabolites in total. Thirteen metabolites were significantly different among the three groups: Rapid ALS, Slow ALS and healthy controls (Fig. 1B and Table. S2). Random forest analysis, whose predictive accuracy for discrimination of three groups was 61% and above random chance, showed that *N*-linoleoyl taurine, a metabolite belonging to the expanded ECS, was the most effective metabolite for distinguishing the groups (Fig.1C). The top-ranked metabolites in the random forest analysis were mainly related to endocannabinoid, nucleic sugar, acylcarnitine, and polyamine metabolism. Similarly, heatmap analysis and sparse partial least squares discriminant analysis (sPLS-DA) also demonstrated that endocannabinoids, nucleic sugar, acylcarnitine, and polyamine metabolites were the discriminant metabolic factors for the three groups (Fig. 1, D to F).

To identify metabolic factors relevant to the disease progression of ALS, we further performed a comparison between the Rapid ALS and Slow ALS groups. Volcano plot and heatmap analyses showed an increase in endocannabinoid and polyamine metabolites in the Rapid ALS group, and pathway enrichment analyses also showed alterations in those metabolic pathways (fig. S1, S2, S3, and S4). In addition, altered levels of bile acid metabolites were also found in the Rapid ALS group with pathway analyses (fig. S3 and table S3). According to these clinical metabolome analyses, we selected five metabolic pathways, endocannabinoid, polyamine/arginine, xanthine, bile acid, and acylcarnitine metabolism, as candidate metabolic biomarkers associated with the progression of ALS.

**Fig. 2.**
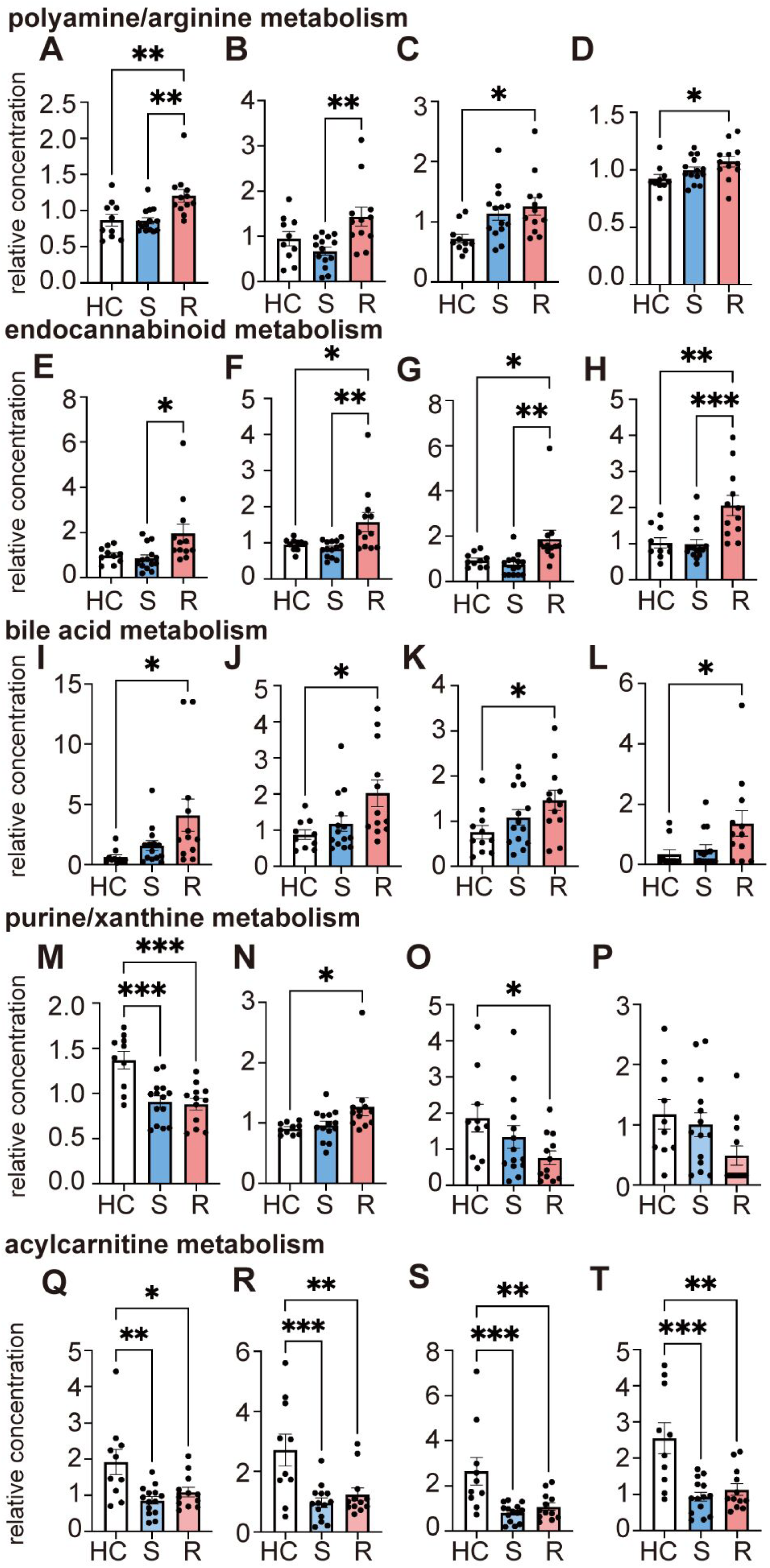
Differences in metabolic pathways among patients with rapidly progressive ALS, patients with slowly progressive ALS, and healthy individuals. Serum levels of representative metabolites in the pathways we focused on were compared among Rapid ALS patients, Slow ALS patients, and healthy individuals. (**A–D**) Polyamine/arginine metabolism (A, (*N*(1) + *N*(8))-acetylspermidine; B, spermidine; C, pro-hydroxy-pro; D, dimethylarginine), (**E–H**) endocannabinoid metabolism (E, *N*-oleoyl taurine; F, *N*-stearoyl taurine; G, *N*-linoleoyl taurine; H, *N*-palmitoyl taurine), (**I–L**) bile acid metabolism (I, cholate; J, hyocholate; K, chenodeoxycholate; L, 7-ketodeoxycholate). (**M–P**) purine/xanthine metabolism (M, adenine; N, xanthine; O, 1,3-demethylurate; P, 3,7-dimethylurate), (**Q–T**) acylcarnitine metabolism (Q, cis-4-decenoylcarnitine; R, 5-dodecenoylcarnitine; S, decanoylcarnitine; T, laurylcarnitine). Data information One-way ANOVA and Tukey’s post hoc analysis were performed (\**p* < 0.05, \*\**p* < 0.01 and \*\*\**p* < 0.001). HC, healthy controls; R, Rapid ALS; S, Slow ALS. Error bars indicate the SEM.

**Fig. 3.**
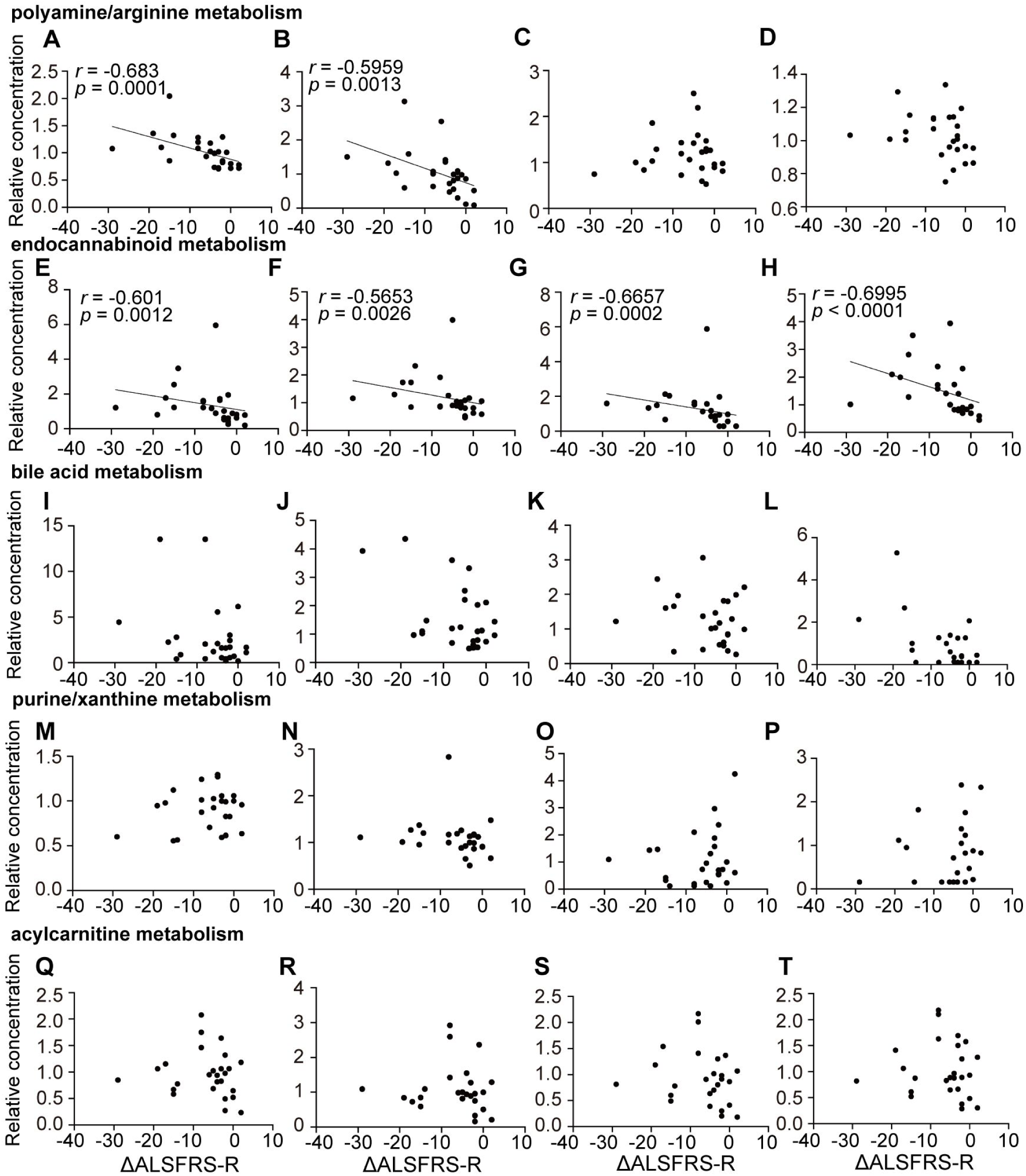
Correlation between serum levels of metabolites and ALSFRS-R decline. We analyzed the correlation between the serum levels of metabolites shown in Figure 2 and the prospective, longitudinal change in ALSFRS-R between the first and second evaluations (ΔALSFRS-R). (**A–D**) Polyamine and arginine metabolism (A, (*N*(1) + *N*(8))-acetylspermidine; B, spermidine; C, pro-hydroxy-pro; D, dimethylarginine), (**E–H**) expanded endocannabinoid metabolism (E, *N*-oleoyl taurine; F, *N*-stearoyl taurine; G, *N*-linoleoyl taurine; H, *N*-palmitoyl taurine), (**I–L**) bile acid metabolism (I, cholate; J, hyocholate; K, chenodeoxycholate; L, 7-ketodeoxycholate), (**M–P**) acylcarnitine metabolism (M, cis-4-decenoylcarnitine; N, 5-dodecenoylcarnitine; O, decanoylcarnitine; P, laurylcarnitine), (**Q–T**) purine/xanthine metabolism (Q, adenine; R, xanthine; S, 1,3-demethylurate; T, 3,7-dimethylurate) Data information Coefficients (*r*) and *p* were shown only for metabolites with strong correlation (*r* > 0.5) in Spearman’s rank correlation coefficient.

We then compared the metabolites belonging to the pathways we focused on among the three groups (Fig. 2 and table S4). The results showed that, in general, the Rapid ALS group had elevated levels of metabolites of the polyamine/arginine, endocannabinoid, and bile acid pathways (Fig. 2, A to L). In contrast, the levels of most metabolites of the purine/xanthine and acylcarnitine pathways were decreased in the ALS groups, particularly in the Rapid ALS group (Fig. 2, M to T). We next investigated the relationship between the serum levels of metabolites and the longitudinal change in motor function (Fig. 3). The results demonstrated that the levels of metabolites belonging to the polyamine/arginine and the expanded ECS, including (*N*(1) + *N*(8))-acetylspermidine and spermidine in the polyamine/arginine pathway and NATs such as *N*-oleoyl taurine, *N*-stearoyl taurine, *N*-linoleoyl taurine, and *N*-palmitoyl taurine in the expanded ECS, were correlated with the change in the ALSFRS-R (Fig. 3, A, B, and E to H). The expanded ECS metabolites associated with the decrease in the ALSFRS-R score were mainly NATs, whereas the other endocannabinoids such as *N*-acyl ethanolamines (NAE) were not altered among the three groups (fig. S5).

### In vitro drug screening

The results of the clinical study on the relationship between the metabolome and disease progression of sporadic ALS patients led us to drug screening using cellular models of ALS. A total of 26 chemical compounds that potentially modulate the focused metabolic pathways (endocannabinoid, polyamine/arginine, xanthine, bile acid, and acylcarnitine), together with riluzole and edaravone as controls, were administered to two cellular models of ALS: NSC-34 cells expressing mutant TDP-43^A315T^ and those expressing mutant SOD1^G93A^ (Fig. 4 and table S5). Among the tested compounds, allantoin and PF-04457845 improved cell viability in both models in a concentration-dependent manner (Fig. 4, B and D). Although arachidonoyl ethanolamide (AEA, anandamide), agmatine, and eflornithine improved the viability of NSC-34 cells expressing mutant TDP-43^A315T^, such efficacy was not verified in the cells expressing mutant SOD1^G93A^ (Fig. 4, E to G).

**Fig. 4.**
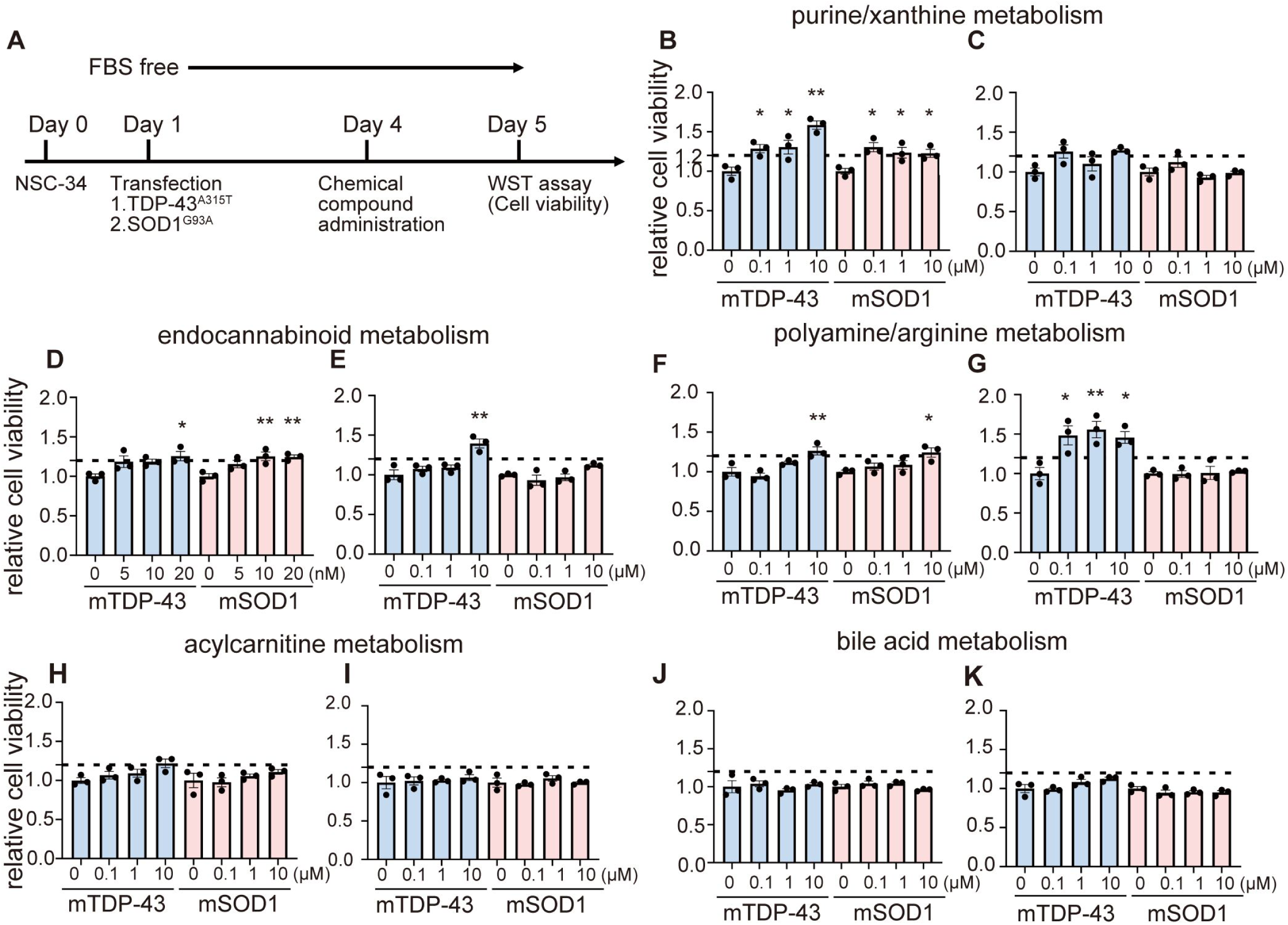
Cellular assay of compounds targeting selected metabolic pathways. We tested a total of 26 compounds that target the identified metabolic pathways. We used NSC-34 cells transfected with TDP-43^A315T^ or SOD1^G93A^ and measured cell viability using WST-8. The results of 10 representative agents are shown in this figure, and the other results are summarized in Supplemental Table 5. (**A**) Protocol of the cell viability assay. (**B–K**) Effects of compounds targeting purine/xanthine metabolism (B, allantoin; C, theobromine), expanded endocannabinoid metabolism (D, PF-04457845; E, arachidonoyl ethanolamide [AEA]), polyamine/arginine metabolism (F, agmatine; G, eflornithine), acylcarnitine metabolism (H, riboflavin; I, clofibrate), and bile acid metabolism (J, glycocholate; K, glycoursodeoxycholate [GUDCA]) on cell viability.Data information One-way ANOVA and Dunnet’s post hoc analysis comparing each concentration to the baseline concentration were performed (\**p* < 0.05 and \*\**p* < 0.01). Error bars indicate the SEM.

Allantoin is a downstream product of the xanthine metabolic pathway that is produced through the metabolism of urate by xanthine oxidase. Similar to urate, allantoin itself has anti-ROS activity (Tzeng *et al*, 2022). PF-04457845 is a fatty acid amide hydrolase (FAAH) inhibitor. FAAH degrades endocannabinoids, including NATs and NAEs, and knockdown of FAAH partially attenuates motor dysfunction in SOD1^G93A^ transgenic mice (Bilsland *et al*, 2006). Based on the results of cellular screening and the putative mechanism of the hit compounds, we further investigated allantoin and PF-04457845 in vivo.

### In vivo analysis of hit compounds in SOD1^G93A^ transgenic mice

The treatment of SOD1^G93A^ transgenic mice with allantoin or PF-04457845 was started at the age of 8 weeks and continued until the ethical endpoint. Body weight measurements and rotarod tests were repeated weekly from the age of 8 weeks to 16 weeks, which was the end stage of this model. Allantoin was mixed with chow at a dose of 0.01% and given to SOD1^G93A^ transgenic mice. The dosage of allantoin was estimated to be ∼10 mg/kg/day based on the average amount of food consumption of 8–16-week-old mice. There was no significant change in their survival between the allantoin-treated and untreated SOD1^G93A^ transgenic mice (mean survival: 132 days in allantoin-treated group, 12 male and 11 female mice; 129.5 days in untreated group, 17 male and 17 female mice, adjusted *p* = 0.2795) (Fig. 5A). PF-04457845 was also mixed with chow at a dose of 0.001%, which corresponded to ∼1 mg/kg/day, and was administered to SOD1^G93A^ transgenic mice. The survival of SOD1^G93A^ transgenic mice administered PF-04457845 was extended by 8.5 days compared with that of untreated mice (mean survival: 138 days in PF-04457845-treated group, 13 males and 15 females; 129.5 days in untreated group, 17 males and 17 females, adjusted *p* = 0.0003) (Fig. 5A). The administration of PF-04457845 also attenuated weight loss and motor functional decline (Fig. 5, B and C).

**Fig. 5.**
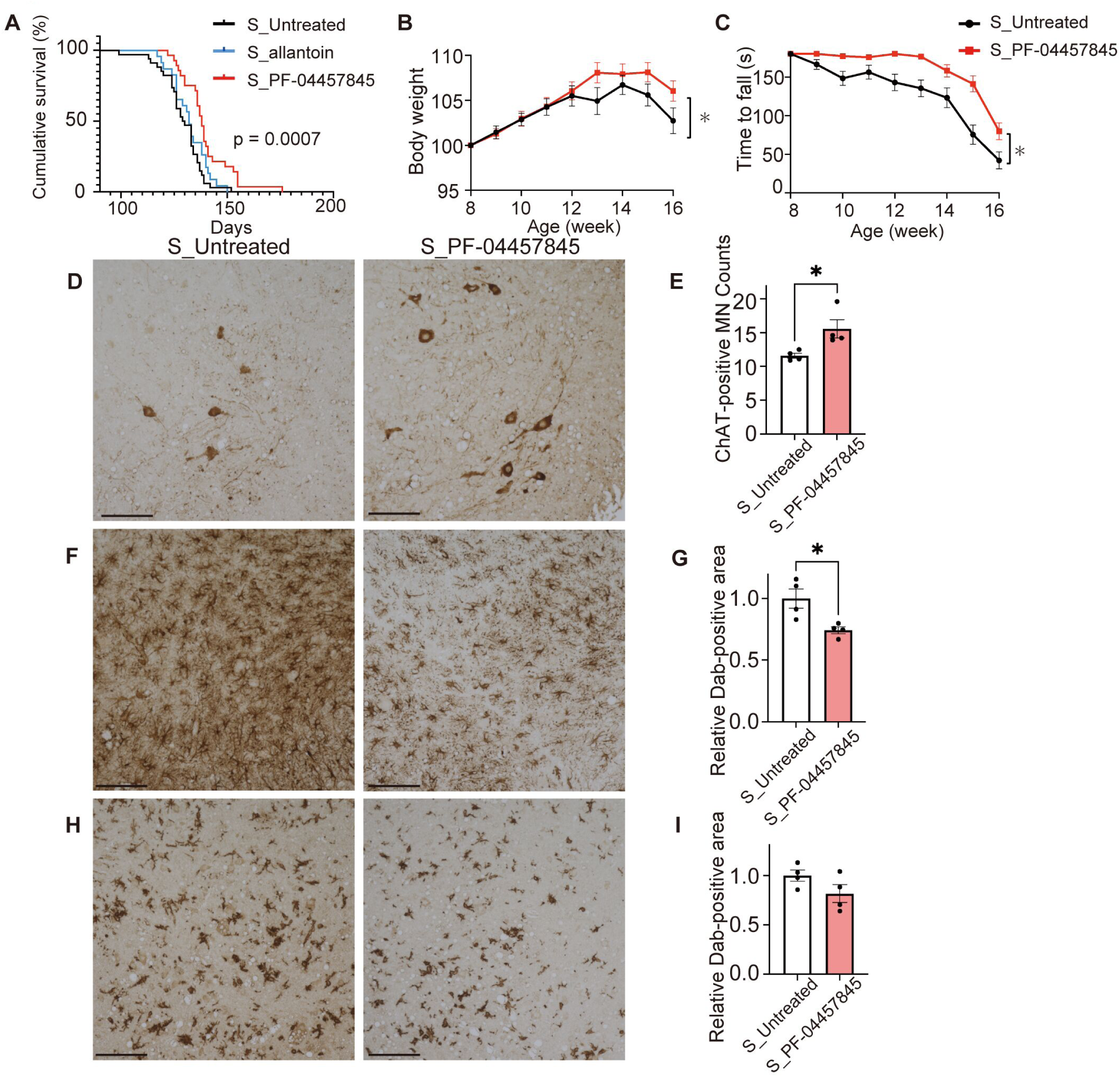
In vivo analysis of hit compounds in SOD1^G93A^ transgenic mice. Allantoin (∼10 mg/kg/day), PF-04457845 (∼1 mg/kg/day), or unmodified chow was orally administered to SOD1^G93A^ transgenic mice from the age of 8 weeks. (**A**) Survival of SOD1^G93A^ transgenic mice treated with allantoin (S_allantoin), PF-04457845 (S_PF-04457845), or unmodified chow (S_Untreated) (n = 23, 28, 34; mean survival, 132, 138, 129.5 days, respectively; *p* = 0.0007, log-rank test for comparison of 3 groups). The administration of PF-04457845 extended the survival of SOD1^G93A^ transgenic mice by 8.5 days compared with that of untreated mice (adjusted *p* = 0.0003, log-rank test with the Holm-Šídák method). Change in body weight (**B**) and rotarod test (**C**) in SOD1^G93A^ transgenic mice treated with PF-04457845 or unmodified chow (\**p* < 0.05 at 16 weeks, as determined by the unpaired *t* test). (**D–I**) Immunohistochemistry images and quantitative analysis of immunoreactivity for choline acetyltransferase (ChAT) (D, E), glial fibrillary acidic protein (GFAP) (F, G), and ionized calcium-binding adapter molecule 1 (IBA-1) (H, I) in the ventral horn of the spinal cords of 16-week-old mice treated with PF-04457845 (n = 4) or unmodified chow (n = 4). Data information Scale bars: 100 µm, Error bars indicate the SEM. \**p* < 0.05, unpaired two-sided *t* test.

To examine the effect of PF-04457845 on the motor neurons of SOD1^G93A^ transgenic mice, paraffin-embedded sections of lumber spinal cord (L5) samples from 16-week-old mice were analyzed by immunohistochemistry. The number of motor neurons with immunoreactivity for choline acetyltransferase within the lumbar spinal cord was significantly preserved in the SOD1^G93A^ mice treated with PF-04457845 compared with untreated mice (Fig. 5, D and E). Proliferations of astrocyte and microglia were evaluated with anti-glial fibrillary acidic protein (GFAP) antibody (Fig. 5, F and G) and anti-ionized calcium-binding adapter molecule 1 (IBA-1) antibody (Fig. 5, H and I), respectively. The proportion of astrocytes was significantly decreased by PF-04457845 treatment, although the proportion of microglia was unchanged.

### Lipidomics analysis of the murine spinal cords

To clarify the influence of PF-04457845 on lipid metabolism, a comprehensive lipidomics analysis was conducted on the spinal cords of three distinct groups: wild-type mice, untreated SOD1^G93A^ transgenic mice, and SOD1^G93A^ transgenic mice treated with PF-04457845. A total of 799 lipid metabolites were identified in the process (table S6). Subsequent sPLS-DA analysis effectively segregated these three groups, underscoring that the differentiation was primarily attributed to the concentrations of NATs and NAEs, both of which are known substrates of FAAH (Fig. 6, A and B). By examining the levels of specific metabolites belonging to NATs and NAEs, we found that both lipids experienced an increase after treatment with PF-04457845 (Fig. 6, C and D). Furthermore, several NATs were found to be elevated in the spinal cord of untreated SOD1^G93A^ transgenic mice compared to wild-type mice, while there was no change in NAEs (Fig. 6, C to E), suggesting a common pathogenic mechanism between SOD1^G93A^ transgenic mice and Rapid ALS patients.

**Fig. 6.**
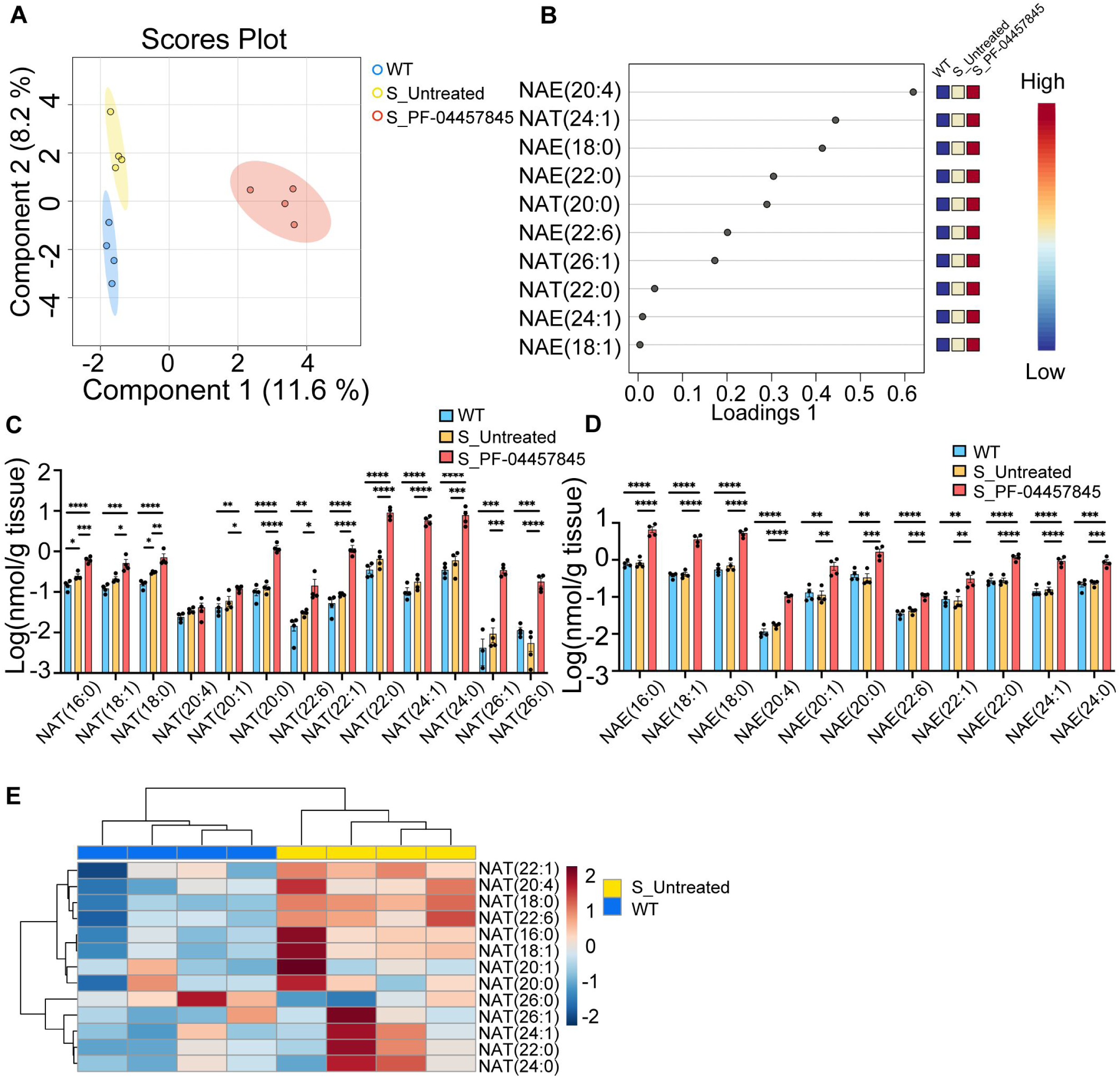
Lipidomics analysis of spinal cords of wild-type and SOD1^G93A^ transgenic mice with or without treatment with PF-04457845. (**A**) Sparse partial least squares discriminant analysis (sPLS-DA) plots by MetaboAnalyst 5.0 to discriminate wild-type mice (WT, n = 4), untreated SOD1^G93A^ transgenic mice (S_Untreated, n = 4) and SOD1^G93A^ transgenic mice treated with PF-04457845 (S_PF-04457845, n = 4). Loading plots of the top 10 lipids by sPLS-DA are shown in (**B**). (**C**) Log transformed concentration of *N*-acyl taurines (NAT). (**D**) Log transformed concentrations of *N*-acyl ethanolamine (NAE). (**E**) Heatmap analysis by *N*-acyl taurines comparing untreated SOD1^G93A^ transgenic mice and wild-type mice. Data information Error bars indicate the SEM. *p < 0.05, **p < 0.01 and ***p < 0.001, ****p < 0.0001. One-way ANOVA and Tukey’s post hoc analysis were performed.

### RNA and protein analysis of the spinal cord of SOD1^G93A^ transgenic mice treated with PF-04457845

To explore the mechanism of phenotypic amelioration in SOD1^G93A^ transgenic mice by PF-04457845, we performed RNA-Seq on their spinal cord. GO analysis of the RNA-Seq data revealed activation of the *Erk1/2* cascade in PF-04457845-treated mice (n = 4) compared with untreated mice (n = 4) (Fig. 7A, table S7). The *Erk1/2* pathway is downstream to the endocannabinoid receptor type 1 (CB1) and type 2 (CB2). CB1 is mainly expressed in neurons and astrocytes, while CB2 is expressed in microglia in the central nervous system (Cristino *et al*, 2020). Ingenuity Pathway Analysis (IPA) analysis of the RNA-Seq data revealed activation of the interleukin-13 (IL-13) pathway in SOD1^G93A^ transgenic mice treated with PF-04457845 (Fig. 7B). In line with this finding, CD36, a downstream molecule of IL-13, was up-regulated in the spinal cord of PF-04457845-treated mice (fig. S6). CD36 is also an M2 microglial marker (Zhao *et al*, 2020; Madathil *et al*, 2018), and IL-13 promotes M1-to-M2 microglial polarization (Orihuela *et al*, 2016). CB2 receptor signaling is also known to activate M1-to-M2 microglial polarization (Mecha *et al*, 2015; Tanaka *et al*, 2020). Therefore, we examined M1 and M2 microglial markers and molecules downstream of CB2 receptor signaling in the mouse spinal cord by Western blotting. The ratio of phosphorylated ERK1/2 to total ERK1/2 was increased in PF-04457845-treated mice compared with untreated mice, suggesting activation of the CB1/CB2 signaling pathway (Fig. 7, C and D). The protein levels of IBA-1 were not significantly changed by PF-04457845 treatment, consistent with the histopathological findings (Fig. 7, C and E). The protein levels of CD86, an M1 microglial marker, were decreased and those of CD36, an M2 marker, were increased in the PF-04457845-treated mice, indicating M1-to-M2 microglial polarization by the treatment (Fig. 7, C, F, and G). The expression of SOD1 was not affected by PF-04457845 (Fig. 7, C and H).

**Fig. 7.**
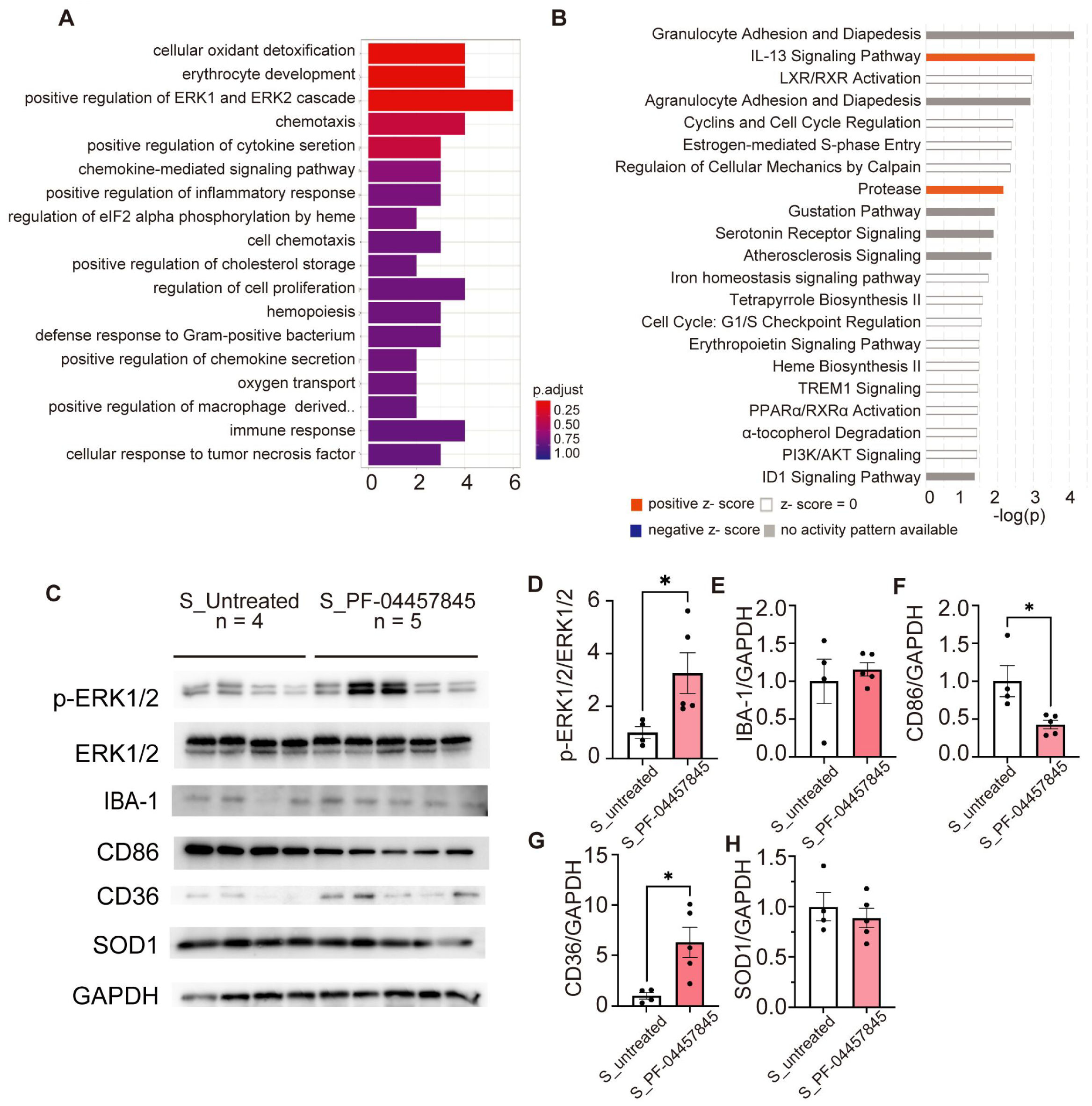
RNA-Seq analysis of the spinal cords from SOD1^G93A^ transgenic mice treated with PF-04457845. (**A–B**) Gene Ontology (GO) enrichment analysis for up-regulated genes (A) and Ingenuity Pathway Analysis (IPA) (B) analysis of genes with significant expression changes (fold change < –1.5 or > 1.5, *p* < 0.05) in the spinal cords of 16-week-old SOD1^G93A^ transgenic mice treated with PF-04457845 (n = 4) compared to those untreated (n = 4). (**C**) Western blot analysis of proteins downstream of cannabinoid receptor type 2 (CB2), microglial markers, and superoxide dismutase 1 (SOD1) in the spinal cords of 16-week-old SOD1^G93A^ transgenic mice treated with PF-04457845 (S_PF-0445785, n = 5) and untreated SOD1^G93A^ transgenic mice (S_untreated, n = 4). (**D–H**) Quantitative analysis of phosphorylated extracellular signal-regulated kinase (ERK)/ERK (D), ionized calcium-binding adapter molecule 1 (IBA-1) (E), CD86 (F), CD36 (G), and SOD1 (H). The levels of IBA-1, CD86, CD36, and SOD were normalized to glyceraldehyde-3-phosphate dehydrogenase (Gapdh) levels. Error bars indicate the SEM. \**p* < 0.05 and \*\**p* < 0.01, unpaired two-sided *t* test.

### Transcriptome alteration in neurons by PF-04457845

The bulk RNA-Seq of the spinal cord primarily revealed altered signaling related to the CB2 receptor, which is mainly expressed in microglia and oligodendrocytes. Given that neurons, as well as astrocytes, express CB1 receptors, the treatment-related changes in neurons did not appear to be reflected in the RNA-Seq data. To examine the impacts of treatment with PF-04457845 on neurons, we performed single-nucleus RNA-Seq (snRNA-Seq) on the spinal cords from SOD1^G93A^ transgenic mice treated with PF-04457845 or unmodified chow (Fig. 8A, table S8). Unsupervised clustering identified six major cell types, and each cluster expressed cell type-specific cell markers (Fig. 8, B and C). The cluster of neurons expressed *Cnr1*, the gene encoding the CB1 receptor; however, *Cnr2*, the CB2 receptor, was barely expressed in the neuronal population (Fig. 8, D and E). The administration of PF-04457845 up-regulated the expression of genes including *Tshz3*, *Ppp3r1*, and *Cyfip1* in neurons (Fig. 8F). Enrichment analysis revealed that PF-04457845 activates synapse plasticity and neuronal development, although no specific pathway was downregulated with the treatment (Fig. 8, G and H).

**Fig. 8.**
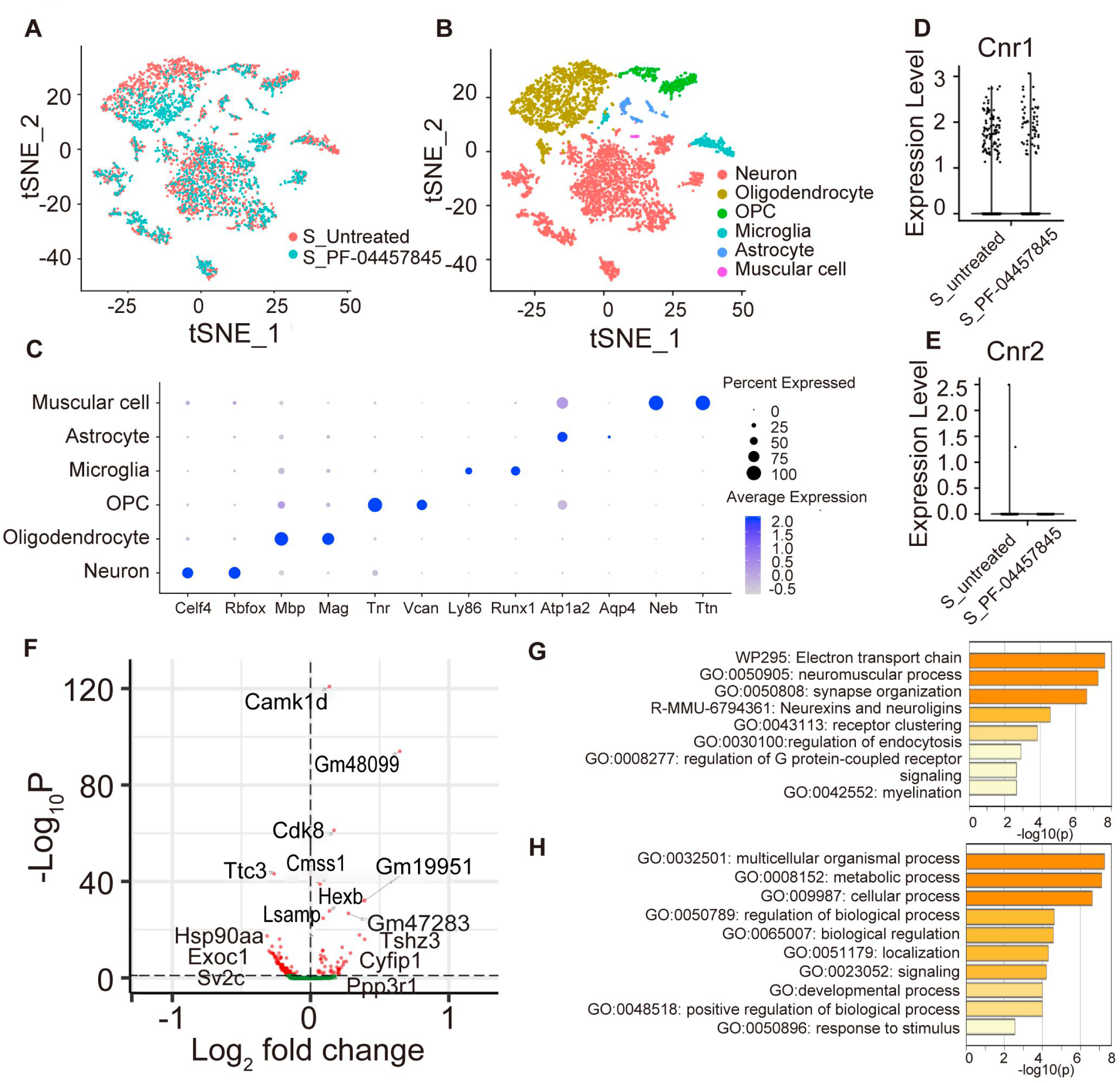
Single-nucleus RNA-Seq analysis of the spinal cords from SOD1^G93A^ transgenic mice treated with PF-04457845. We performed single-nucleus RNA sequencing on the spinal cords of 16-week-old SOD1^G93A^ transgenic mice treated with PF-04457845 (S_PF-04457845) or unmodified chow (S_Untreated). (**A**) t-Distributed stochastic neighbor embedding (t-SNE) plot color-coded by sample, with red for untreated mice and blue for PF-04457845-treated mice. (**B**) t-SNE plots of all cells sequenced showing six cell types. (**C**) Dot plot depicting the expression of specific markers for each cell type. (**D–E**) Expression levels of CB1 (D) and CB2 (E) in the neuron cluster of each sample. (**F**) Volcano plot of the differentially expressed genes between the neuron cluster from the mice treated with PF-04457845 compared to that from the untreated mice. (**G–H**) Gene Ontology (GO) enrichment analysis for up-regulated (G) and downregulated (H) genes between the neuron clusters from the mice treated with PF-04457845 and the untreated mice.

## DISCUSSION

In the present study, metabolome analysis of sera identified metabolic pathways associated with the progression rates of sporadic ALS. Investigation of individual metabolites showed that the serum levels of expanded ECS metabolites, especially NATs, were correlated with longitudinal changes in the ALSFRS-R. Based on these clinical findings, we performed an in vitro screening of compounds that target the identified metabolic pathways. Among the hit compounds in the cellular assay, PF-04457845, an FAAH inhibitor which up-regulates NATs and NAEs, ameliorated motor neuron degeneration in SOD1^G93A^ transgenic mice, presumably through the regulation of microglial and neuronal function.

Based on the data analysis of sporadic ALS patient serum metabolomics, we selected the following five key metabolic pathways that showed differences between rapidly and slowly progressive groups: endocannabinoid, purine/xanthine, polyamine/arginine, bile acid, and acylcarnitine metabolism. Previous studies on metabolomics in ALS analyzed the plasma or cerebrospinal fluid (CSF) of patients and healthy individuals, although the results vary due to differences in samples and metabolomic methodologies (Blasco *et al*, 2016, 2010, 2013, 2014; Veyrat-durebex *et al*, 2019; Patin *et al*, 2017; Lee *et al*, 2021; Goutman *et al*, 2022, 2020). The present study partially reproduced the results of such previous studies. For instance, we identified altered serum levels of adenine, xanthosine, and xanthine in ALS patients. Alterations in purine/xanthine pathways have also been reported in previous studies (Veyrat-durebex *et al*, 2019; Lee *et al*, 2021), suggesting that reactive oxygen species (ROS) generated by xanthine oxidase are related to ALS etiology. Allantoin, a hit compound of our in vitro screening, is a catabolic product of uric acid that has anti-ROS activity (Tzeng *et al*, 2022; Papandreou *et al*, 2019). In addition, aldioxa, an anti-peptic ulcer drug containing allantoin, showed therapeutic effects on iPSC-derived motor neurons from sporadic ALS patients (Fujimori *et al*, 2018). We thus tested allantoin in SOD1^G93A^ transgenic mice, but the compound did not show a detectable benefit in the present study. The reasons for this failure may include limited antioxidative properties in vivo and an insufficient concentration in the central nervous system. Our study also demonstrated alterations in the levels of acylcarnitine, a metabolite involved in β-oxidation of fatty acids in mitochondria, which has been reported in previous metabolomic studies (Goutman *et al*, 2022, 2020).

The major discovery of the present study is that markers of the expanded ECS are up-regulated in sporadic ALS patients with rapid progression. In particular, serum levels of NATs were increased in the rapidly progressive ALS group, being consistent with the result of lipidomics analysis on the spinal cord of SOD1^G93A^ transgenic mice. Furthermore, the serum levels of NATs were closely correlated with the longitudinal decrease in the ALSFRS-R rating, while major endocannabinoids, AEA and 2-AG, were unchanged in subjects with ALS. Although functions of NATs in the central nervous system has yet to be elucidated, our results suggest that NATs are aberrantly regulated in patients and the mouse model of ALS and associated with disease progression in sporadic ALS.

In our cellular experiments, neuroprotective effects of PF-04457845, an inhibitor of FAAH, are stronger than those of AEA or 2-AG. Moreover, pharmacological inhibition of FAAH led to a substantial increase in the levels of both NATs and NAEs in the spial cord, resulting in phenotypic amelioration of the mutant SOD1 mice. Collectively, targeting the expanded ECS, including NATs and NAEs, appears to be a common therapeutic approach for both human patients and the animal model of ALS. Our findings also suggest that the elevated levels of NATs in rapidly progressive ALS are indicative of defensive metabolic responses, and that further up-regulation of such compensative machinery leads to mitigation of disease progression. PF-04457845 is known to increase the concentration of endocannabinoids in tissues, including those in the central nervous system (Ahn *et al*, 2011), and has been examined in a phase 2 clinical trial for pain associated with osteoarthritis, which demonstrated the safety and tolerance of this compound in human (Huggins *et al*, 2012). Taken together, our results suggest that PF-04457845 is a novel candidate of disease modifying therapy for ALS.

Major molecules of the ECS act through endocannabinoid receptors CB1 and CB2. For instance, NAEs including AEA, as well as 2-AG, bind to CB1 and CB2 as well as to transient receptor potential cation channel subfamily V member 1 (TRPV1), while NATs are known as ligands for TRPV1 and TRPV4. In previous studies, activation of the CB1 and CB2 pathways demonstrated potential benefits in SOD1^G93A^ mice (Bilsland *et al*, 2006; Pasquarelli *et al*, 2017). The bulk RNA-Seq data on the spinal cords from treated mice revealed activation of the IL-13 and CB2 signaling pathways, both of which are known to facilitate microglial polarization from the M1-to M2-dominant state (Orihuela *et al*, 2016; Cristino *et al*, 2020). It is also known that the activation of TRPV1 brings M2-dominant microglial polarization (Bok *et al*, 2018). Given that M1 and M2 microglia populations have proinflammatory and anti-inflammatory properties, respectively, the effect of PF-04457845 in our study is, at least partially, attributable to M2 microglial polarization. In addition, our study also demonstrated the benefits of PF-04457845 in NSC-34 cells expressing pathogenic TDP-43 or SOD1 protein, suggesting the effect of this compound on neurons. In support of this view, the results of snRNA-Seq indicated that *Tshz3*, *Cyfip1*, and *Ppp3r1* are up-regulated in neurons treated with PF-04457845 in vivo. Tshz3 is essential for cerebral cortical projection neuron development and has been implicated in the pathogenesis of autism spectrum disorder (ASD) (Caubit *et al*, 2016). Cyfip1 is also related to ASD and schizophrenia and is known to play an important role in dendritic spine maturation, synaptic activities, and plasticity and acts with CB1 (Monday *et al*, 2020; Cioni *et al*, 2018; Hsiao *et al*, 2016; DeRubeis *et al*, 2013; Davenport *et al*, 2019). Ppp3r1 is a calcineurin subunit that is required for neuronal survival and neurite outgrowth of mesencephalic dopaminergic neurons by glial cell line-derived neurotrophic factor (Consales *et al*, 2007). Variants in *PPP3R1* are associated with the accelerated progression of Alzheimer’s disease(Peterson *et al*, 2014). Altogether, in addition to the microglial mechanism mentioned above, PF-04457845 appears to have direct protective effects on neurons via CB1-mediated pathways.

Another finding of the present study is an altered polyamine/arginine metabolism in sporadic ALS patients. Our results showed that serum levels of dimethylarginine, spermidine, and (*N*(1) + *N*(8))-acetylspermidine were increased in the rapidly progressive ALS group, and the levels of spermidine and (*N*(1) + *N*(8))-acetylspermidine were well correlated with the change in ALSFRS-R. A previous report exploring human plasma and SOD^G93A^ transgenic mice plasma, cerebral cortex, and muscle tissue by metabolomics identified alterations in arginine and polyamine metabolism as common metabolic phenotypes of human ALS and SOD1^G93A^ transgenic mice mouse (Patin *et al*, 2017). Elevated levels of asymmetrical dimethylarginine in CSF are shown to be a poor prognostic factor of ALS (Ikenaka *et al*, 2019). In addition, alterations of acylcarnitine, xanthine, and polyamine were also observed in Parkinson’s disease, suggesting a common metabolic dysregulation in neurodegeneration regardless of causative proteins (Hatano *et al*, 2016; Saiki *et al*, 2017, 2019). Given that the polyamine pathway is related to the cell cycle and autophagy(Saiki *et al*, 2019), the dysregulation of polyamine metabolites may lead to neuronal cell death and muscle atrophy. In our in vitro screening, agmatine and eflornithine, compounds related to polyamine/arginine metabolism, improved the viability of NSC-34 cells expressing pathogenic TDP-43 but were not effective in the SOD1^G93A^ cell model.

The present study has several limitations. First, the sample size of the clinical study was small. Therefore, larger validation studies are needed. Second, we utilized two murine cellular models for in vitro screening and SOD1^G93A^ transgenic mice for in vivo analysis to analyze motor neurons and glial cells. These models do not necessarily reflect the pathogenesis of sporadic ALS, though our study demonstrated the common elevation of NATs in rapid progressive ALS patients and the murine model. Assays using human cells including iPSC-derived motor neurons may strengthen the results of the present study. Finally, the efficacy of the FAAH inhibitor PF-04457845 seems superior to similar approaches in the literature, but the pharmacological basis for the difference remains unclear.

## METHODS

### Participants

Subjects who were clinically diagnosed with the revised El Escorial criteria of definite, probable, or possible ALS were consecutively recruited. The principal inclusion criteria were no family history and disease duration of ≤ 2 years at the time of enrollment and a mini mental state examination score of ≥ 23. Subjects who had severe complications such as malignancy, heart failure, or renal failure were excluded from this study. Subjects with sporadic ALS were assessed during hospitalization at the initial evaluation and follow up evaluations at an outpatient clinic were conducted every 6 months. Age– and sex-matched healthy controls were also recruited during the same period as the ALS patients. All study subjects were Japanese and observed at the Nagoya University Hospital between May 2013 and December 2017.

### Clinical evaluation and sample collection

Disease onset was defined as the time point when the subject felt weakness of any body part. Patients with ALS were classified according to onset type and were sorted into either the limb-onset type group or the bulbar-onset type group, according to the site where they first felt weakness. Disease severity was assessed with the Japanese version of the revised ALS Functional Rating Scale (ALSFRS-R), a validated questionnaire-based functional rating scale for ALS (Ohashi *et al*, 2001).

At the first evaluation, venous blood samples were collected from ALS patients in the supine position after more than 12 hours of fasting and just after waking up during hospitalization. At the outpatient clinic, venous blood samples of healthy individuals in the sitting or supine position were collected after more than 12 hours of fasting. Serum samples were centrifuged at 3000 rpm for 10 min and stored at –80 °C until processing at Metabolon Inc.

### Metabolomics analysis

Untargeted metabolomics profiling of serum samples was performed using ultrahigh-performance liquid chromatography-tandem mass spectrometry (UPLC‒MS/MS) by Metabolon (Durham, NC). Several recovery standards were added prior to the first step in the extraction process for QC purposes. To remove protein, dissociate small molecules bound to protein or trapped in the precipitated protein matrix and to recover chemically diverse metabolites, proteins were precipitated with methanol. The resulting extract was divided for analysis by two separate reversed-phase (RP)/UPLC‒MS/MS methods with positive ion mode electrospray ionization (ESI), for analysis by RP/UPLC‒MS/MS with negative ion mode ESI, and for analysis by HILIC/UPLC‒MS/MS with negative ion mode ESI. In addition to the spiked internal standards within each sample, a pooled ‘technical replicate’ generated from all study samples was periodically injected into the UPLC‒MS/MS to assess instrument performance and calculate overall process and platform variability. Compounds were identified by comparison to the library entries of purified standards or recurrent unknown entities, and relative levels were quantitated. Missing values were replaced by the minimal value determined for that metabolite in the entire cohort, per Metabolon protocols.

### Metabolomics data analysis

Volcano plots, one-way ANOVA, heatmap analysis, and enrichment analysis were performed using MetaboAnalyst 5.0 (Pang *et al*, 2021). For multivariate analysis, sparse partial least squares discriminant analysis (sPLS-DA) was performed to discriminate between rapid ALS patients, slow ALS patients and healthy individuals using MetaboAnalyst 5.0. Loading plots indicated the importance of metabolites for sPLS-DA. Adding to this multivariate analysis, random forest (RF) classification in R software was applied to determine whether the differences in the identified metabolites effectively separated the three groups. The mean decrease in accuracy, which represents variable importance, was calculated.

### Cell culture and transfection

Mouse NSC-34 motor neuron-like cells (kindly provided by N.R. Cashman, University of British Columbia, Vancouver, Canada) were cultured in a humidified atmosphere of 95% air/5% CO2 in a 37 °C incubator. NSC-34 cells were maintained in Dulbecco’s Modified Eagle’s Medium (DMEM) containing 10% (v/v) fetal bovine serum (Life Technologies) with 5% penicillin/streptomycin (Wako). The plasmids were transfected by using OPTI-MEM (Gibco) and Lipofectamine 2000 (Invitrogen) according to the manufacturer’s instructions. NSC-34 cells were differentiated in DMEM for 48 hours. We prepared the mutant TDP-43^A315T^ and the mutant SOD1^G93A^ vectors as previously described (Niwa *et al*, 2007; Iguchi *et al*, 2012).

### Chemical compounds

PF-04457845 was supplied by Mitsubishi-Tanabe Pharma, and the other compounds were commercially purchased from Cayman Chemical (arvanil); Kanto Chemical (xanthine); MedChemExpress (eflornithine); Focus Bioscience (Mitoquazone); Sigma‒Aldrich (allantoin, arachidonoyl ethanolamide, clofibrate, febuxostat, inosine, metformin, MDL72527, oleic acid, riboflavin, S-adenosyl-L-methionine, Spermidine, Spermine, and 2-arachidonoyl glycerol); TCI, (agmatine, glicocholic acid, glycoursodeoxycholic acid, riluzole, theobromine, theophiline, trimetazidine, and xanthosine), or Wako (arginine and ornithine).

### Cell viability

Cell viability assays were performed using a WST-8 kit (Roche Diagnostics) according to the manufacturer’s instructions. Transfected and differentiated NSC-34 cells were cultured in 96-well plates, and chemical compounds were administered. Twenty-four hours after treatment with the selected chemicals, cells were incubated with the WST-8 substrate for 30–60 min, after which the absorbance of the wells was measured at 450 nm using a plate reader (EnSpireTM, PerkinElmer).

### Transgenic mice

Transgenic mice overexpressing the human SOD1 gene carrying the G93A mutation were purchased from the Jackson Laboratory and maintained as hemizygotes by mating transgenic males with B6/SJL F1 females. For oral administration of allantoin to mice, allantoin was mixed with powdered rodent chow at a concentration of 0.01%, which is consistent with the dosage of 10 mg/kg/day calculated based on the weekly consumption of chow from the age of 8 weeks to 16 weeks. For oral administration of PF-04457845, PF-04457845 was mixed with powdered rodent chow at a concentration of 0.001%, which is consistent with the dose of 1 mg/kg/day. All mice had unlimited access to food and water. Allantoin (10 mg/kg/day) and PF-04457845 (1 mg/kg/day) were administered from week 8 until the ethical endpoint, which was defined as loss of the ability to right itself after being placed on its side or inability to ambulate from a given location.

### Behavioral analysis

All behavioral tests were performed weekly, and the data were prospectively analyzed. The animal rotarod performances were assessed weekly using an Economex Rotarod (Ugo Basile).

### Aminal sample collection

Mice were deeply anesthetized, and the entire spinal cord was dissected and snap-frozen in powdered CO_2_ in acetone for Western-blot and RNA-Seq and by liquid nitrogen for lipidome analysis. The mice were sacrificed at the age of 16 weeks. Mouse tissues were dissected, postfixed with 10% phosphate-buffered formalin and paraffin embedded for histological analysis.

### Immunoblotting

Mouse tissues were lysed with RIPA buffer (50 mM 1 M Tris pH 7.4, 150 mM NaCl, 1% Triton X-100, 1% deoxycholate, 0.1% SDS) using a dounce homogenizer for mouse tissue, sonicated, and centrifuged at 20,000 × g for 15 min at 4 °C. The supernatant was kept as the RIPA soluble fraction, and equal amounts of protein were separated on 5–20% SDS‒PAGE gels (Wako) and transferred to Hybond-P membranes (GE Healthcare). The following primary antibodies and dilutions were utilized: SOD1 (ab51254, 1:1000; Abcam), IBA-1 (016-20001, 1:1000; Wako), CD36 (ab252923, 1:1000; Abcam), CD86 (#1858-1, 1:1000; Epitomics), p44/42 MAPK (#9102, 1:5000; CST), and phosphorylated p44/42 MAPK (#5726, 1:1000; CST). Primary antibodies bound to the proteins were probed with a 1:5000 dilution of horseradish peroxidase-conjugated secondary antibodies, and the bands were detected using an immunoreaction enhancing solution (Can Get Signal; Toyobo) and enhanced chemiluminescence (ECL Prime; GE Healthcare).

Chemiluminescence signals were digitized using a LAS-3000 imaging system (Fujifilm). The signal intensities of independent blots were quantified using ImageJ. Membranes were reprobed with an anti-GAPDH antibody (MAB374, 1:5000; Santa Cruz Biotechnology) for normalization.

### Histology and immunohistochemistry

The lumbar spinal cords were collected from mice. The samples were embedded in paraffin, and 3-µm sections were prepared. The sections designated to be incubated with the anti-ChAT, GFAP, and IBA-1 antibodies were boiled in 10 mM citrate buffer for 15 min. Then, the sections were incubated with a secondary antibody labeled with a polymer as part of the Envision+ system containing horseradish peroxidase (Dako Cytomation). The following primary antibodies and dilutions were used to stain mouse tissues: ChAT (AB144P, 1:100; Millipore), GFAP (#ab53554, 1:1000; Abcam), and IBA-1 (013-27691/1:1000; Wako).

ChAT-immunoreactive neurons in the ventral horn of the lumbar spinal cord were counted in every fifth section from the 50 consecutive sections by using BZ-X810 (Keyence), and the mean total number of ChAT-immunoreactive neurons was compared between treatment groups. The area of ChAT-immunoreactive neurons was analyzed using NIH ImageJ software. GFAP– and IBA-1-positive areas were also calculated by ImageJ software.

### Lipidomics analysis of murine spinal cords

Frozen tissues (about 10 mg) were homogenized with a probe sonicator in methanol/water = 2/0.7 (v/v), and lipids were extracted using the method of Bligh and Dyer with internal standards. The organic (lower) phase was transferred to a clean vial and dried under a nitrogen stream. The lipids were resolubilized in methanol/isopropanol/chloroform = 45/45/10 (v/v/v) and stored at – 80 °C. A portion of the extracted lipids was injected into an ultrahigh-performance liquid chromatography (LC)-electrospray ionization (ESI)–tandem mass spectrometry system (LC-MS/MS). The quantification of FFA was carried out as described previously (Lee-Okada *et al*, 2021). For the quantification of Sulfatide, S1P, NAT, NAEP, LPA, LPG, LPI, LPS, PA, PS, and PT, LC separation was performed on an ACQUITY Premier BEH C18 column (1.7 µm, 2.1 × 50 mm; Waters). Mobile phase A was H_2_O/methanol = 95/5 (v/v%), mobile phase B was isopropanol/methanol = 63/37 (v/v%), and mobile phase C was H_2_O/methanol/28% NH_4_OH = 93/5/2 (v/v/v%). The LC method consisted of a linear gradient from A/C = 95/5 (v/v%) to B/C = 95/5 (v/v%) over 15 min, B/C = 95/5 (v/v%) for 8 min, and equilibration with A/C = 95/5 (v/v%) for 5 min (28 min total run time). The flow rate was 0.3 mL/min, and the column temperature was 25 °C. For the quantification of BMP, an isocratic LC separation was performed with methanol containing 10 mM ammonium formate on a COSMOCORE 2.6C18 column (2.1 × 100 mm; Nacalai Tesque) coupled to an ACQUITY UPLC BEH C18 VanGuard Pre-column (1.7 µm, 2.1 × 5 mm; Waters, Milford, MA, USA). The flow rate was 0.3 mL/min, and the column temperature was 55 °C. For the quantification of other lipid classes, LC separation was performed on an ACQUITY UPLC BEH C18 column (1.7 µm, 2.1 × 100 mm; Waters) coupled to an ACQUITY UPLC BEH C18 VanGuard Pre-column (1.7 µm, 2.1 × 5 mm; Waters). Mobile phase A was acetonitrile/water = 60/40 (v/v%) containing 10 mM ammonium formate and 0.1% (v/v) formic acid, and mobile phase B was isopropanol/acetonitrile = 90/10 (v/v%) containing 10 mM ammonium formate and 0.1% (v/v) formic acid. The LC gradient consisted of 20% B for 2 min, a linear gradient to 60% B over 4 min, a linear gradient to 100% B over 16 min, and equilibration with 20% B for 5 min (27 min total run time). The flow rate was 0.3 mL/min, and the column temperature was 55 °C. Multiple reaction monitoring (MRM) was performed using a Xevo TQ-S micro triple quadrupole mass spectrometry system (Waters) equipped with an ESI source. The ESI capillary voltage was set at 1.0 kV, and the sampling cone was set at 30 V. The source temperature was 150 °C, the desolvation temperature was 500 °C, and the desolvation gas flow was 1,000 L/h. The cone gas flow was 50 L/h.

### Lipidomics data analysis

Lipidomics data of murine spinal cords were analyzed using Metaboanalyst 5.0. For multivariate analysis, sparse partial least squares discriminant analysis (sPLS-DA) was performed to discriminate between wild-type mice, untreated SOD1^G93A^ transgenic mice and SOD1^G93A^ transgenic mice treated with PF-04457845. Loading plots indicated the importance of metabolites for sPLS-DA. The concentrations of N-acyl taurines and N-acyl ethanolamine were log-transformed for One way ANOVA and Tukey’s post-hoc analysis. Heatmap analysis for comparing wild-type mice and untreated SOD1^G93A^ transgenic mice was also performed using Metaboanalyst 5.0.

### RNA-Seq of murine spinal cords

Total RNA was extracted from the thoracic spinal cord using the Invitrogen^TM^ TRIzol^TM^ Plus RNA Purification Kit. The total RNA concentration was calculated by Quant-IT RiboGreen (Invitrogen). To assess the integrity of the total RNA, samples were run on the TapeStation RNA screentape (Agilent). Only high-quality RNA preparations with an RIN greater than 7.0 were used for RNA library construction. A library was independently prepared with 1 µg of total RNA for each sample by an Illumina TruSeq Stranded mRNA Sample Prep Kit (Illumina, Inc.). The first step in the workflow involved purifying the poly-A-containing mRNA molecules using poly-T-attached magnetic beads. Following purification, the mRNA was fragmented into small pieces using divalent cations under elevated temperature. The cleaved RNA fragments were copied into first strand cDNA using SuperScript II reverse transcriptase (Invitrogen) and random primers. This process was followed by second strand cDNA synthesis using DNA Polymerase I, RNase H, and dUTP. These cDNA fragments then went through an end repair process, the addition of a single ‘A’ base, and then ligation of the adapters. The products were then purified and amplified with PCR to create the final cDNA library. The libraries were quantified using KAPA Library Quantification kits for Illumina Sequencing platforms according to the qPCR Quantification Protocol Guide (KAPA BIOSYSTEMS) and qualified using the TapeStation D1000 ScreenTape (Agilent Technologies). Indexed libraries were then submitted to an Illumina NovaSeq (Illumina, Inc.), and paired-end (2 × 100 bp) sequencing was performed by Macrogen Incorporated.

### RNA-Seq data analysis

The quality of the raw paired-end sequence reads was assessed with Fast QC (Version 0.11.5; https://www.bioinformatics.babraham.ac.uk/projects/fastqc/). Low-quality (< 20) bases and adapter sequences were trimmed by Trimmomatic software (Version 0.38) with the following parameters: ILLUMINACLIP: path/to/adapter.fa:2:30:10 LEADING:20 TRAILING:20 SLIDINGWINDOW:4:15 MINLEN:36. The trimmed reads were aligned to the reference genome using the RNA-Seq aligner HISAT2 (Version 2.1.0). The HISAT2-resultant .sam files were converted into .bam files with samtools and used to estimate the abundance of uniquely mapped reads with featureCounts (Version 1.6.3). The raw counts were normalized to transcripts per million (TPM). Differentially expressed genes (DEGs) were identified using DESeq2 (Version 1.24.0) with the threshold of FC (Fold Change) > 1.5 and *p* value < 0.05. Gene set enrichment analysis was conducted by DAVID (Version 1.22.0), and Gene Ontology (GO) terms with *p*.adjust < 0.05 by the Benjamini and Hochberg (BH) method were extracted. DEGs were analyzed by Ingenuity Pathway Analysis (IPA) (Version 51963813; QIAGEN Inc., https://www.qiagenbioinformatics.com/products/ingenuity-pathway-analysis).

### Single-nucleus RNA-Seq of murine spinal cords

#### Nuclei isolation

Spinal cords from 4 mice for each condition were pooled for homogenization and nuclear isolation. Briefly, fresh frozen murine spinal cord was placed in a Dounce homogenizer (Kimble Chase 2 ml Tissue Grinder) containing ice-cold lysis buffer (10 mM Tris-HCl pH 7.4, 10 mM NaCl, 3 mM MgCl2, 0.1% Igepal and 0.2 U/μl RNase inhibitor). After homogenizing with 5 strokes of pestle A and 5–10 strokes of pestle B, the lysate was diluted with 1.4 mL of lysis buffer and incubated for 5 minutes. The homogenate was passed over a 70 μm strainer (Fisher Scientific). The filtered lysate was centrifuged at 500 × g for 5 min at 4 °C. After centrifugation, the pellet was resuspended in 1.5 ml lysis buffer and incubated for 3 minutes on ice. Then, the pellet was centrifuged again at 500 × g for 5 minutes and resuspended in 1500 µl of wash buffer (1× PBS with 1% BSA and 0.2 U/µl SUPERase-In^TM^ RNase Inhibitor (Thermo Fisher). The nuclei were washed and centrifuged at 500 × g for 5 minutes three times and filtered through a 40 µm cell strainer. The cell suspensions were processed according to the debris removal solution (Milteni Biotec) protocol, a density gradient method to remove dead cells and debris. Cells were resuspended in cold PBS with 0.04% BSA and stained with DAPI. The nuclei were sorted with a FACS Aria instrument (Becton Dickson) with a 100 nm nozzle and 405 nm excitation laser. The instrument was controlled by a PC running FACS DiVa™ software (Becton Dickson). These nuclei were centrifuged, inspected for appearance, and concentrated for the next step.

#### Single-nucleus RNA sequencing

The nuclei per sample were run on the 10 × Chromium Single cell 3’ gene expression v3.1 platform. Sequencing libraries were constructed according to the manufacturer’s instructions, and cDNA samples were run on an Agilent Bioanalyzer using the High Sensitivity DNA Chip as quality control and determination of cDNA concentrations. The samples were run on an Illumina HiSeq2500 at Read 1 = 28 bp and Read 2 = 90 bp, with a minimum depth of 20,000 reads per nucleus. For alignment, introns and exons were included in the reference genome (mm10) using the CellRanger v.6.1.2 pipeline (10X Genomics). Sequencing data were analyzed using the R package Seurat Version 4.1.3. The gene barcode matrices for each sample were imported into R using the Read10X function.

#### Quality check analysis

All 10 x runs for each sample were filtered for nuclei with less than 1% contamination of mitochondrial genes, with 200 to 2000 genes per cell, and genes with a count of 1 in at least 3 cells were retained. A total of 4,869 nuclei passed quality control filtering and proceeded to analysis. UMI counts were then normalized in Seurat 3.0, and the top 2000 highly variable genes were identified using the FindVariableFeatures function with variance stabilization transformation (VST).

#### Clustering and data analysis

Principal component analysis (PCA) was performed using the top 2000 variable genes prior to clustering. To visualize profiles in two-dimensional space, t-distributed stochastic neighbor embedding (t-SNE) was performed with the top 25 principal components based on ElbowPlot. Clustering was performed using the FindNeighbors and FindClusters functions in the Seurat R package, and the resolution was 0.15. To identify each cell type, ‘Celf4’ and ‘Rbfox’ were used as specific cell markers for neurons, ‘Mbp” and ‘Mag” for oligodendrocytes, ‘Tnr’ and ‘Vcan” for oligodendrocyte precursor cells (OPCs), ‘Ly86’ and ‘Runx1’ for microglia, ‘Alp1a2’ and ‘Aqp4’ for astrocytes, and ‘Neb’ and ‘Ttn’ for muscular cells. These markers were used to assign cell-type annotations manually for each cell cluster.

#### Single-nucleus RNA-seq data analysis

DEGs between PF-04457845-treated and untreated groups were identified using the FindMarkers function of the Seurat package in R, which functions based on the Wilcox method. Averaged log2 (fold change) of gene expression, the percentage of cells expressing the genes in each group (pct.1 and pct.2), *p* value and adjusted *p* value were generated. DEG lists were produced by filtering all genes for log2-fold changes > 0.1 and adjusted *p* < 0.05. GO terms were extracted as stated above. DEGs up-regulated in PF-04457845-treated mice compared to untreated mice were analyzed with the Metascape portal (www.metascape.org)(Zhou *et al*, 2019).

## Study approval

The clinical part of this study was conducted according to the Declaration of Helsinki, the Ethical Guidelines for Medical and Health Research Involving Human Subjects endorsed by the Japanese government. It was approved by the Ethics Review Committee of Nagoya University Graduate School of Medicine (Nos. 2013-0035 and 2015-0041), and all participants gave written informed consent before participation.

All animal experiments were performed in accordance with the National Institutes of Health Guide for the Care and Use of Laboratory Animals and under the approval of the Nagoya University Animal Experiment Committee (No. 29170).

## Statistical analysis

We analyzed the data by using two-sided unpaired *t* tests and chi-square tests for comparisons of two groups and analysis of variance (ANOVA) for multiple comparisons using GraphPad Prism 9. For multiple comparisons, we utilized Tukey’s test for comparing metabolites among Rapid ALS patients, Slow ALS patients, and healthy individuals and Dunnett’s test for comparing each concentration of chemical compounds and drug screening in ALS cell models. A Spearman rank correlation test was performed to analyze the correlation between the longitudinal differences in ALSFRS-R and serum metabolite levels. In addition, the survival rate was analyzed using Kaplan‒Meier and log-rank tests for comparison of vehicle-treated, allantoin-treated and PF-04457845-treated SOD1^G93A^ transgenic mice. Post hoc analyses for multiple comparisons of each pair of groups and adjusted *p* values were performed with the Holm-Šídák method. We considered *p* values of < 0.05 to indicate statistical significance, and correlation coefficients (*r*) greater than 0.5 were considered to be strong. For experiments using mice, the number of animals is stated in the figure legends.

## Data availability

The metabolomics source was deposited into Mendeley (doi: 10.17632/f8gtkwfs6s.1). The scRNA-Seq and bulk RNA-Seq raw data files were deposited into the NCBI Gene Expression Omnibus (GEO) (https://www.ncbi.nlm.nih.gov/geo/query/acc.cgi?acc=GSE242942/; accession no.GSE242942).

## Acknowledgements Funding

Japan Society for the Promotion of Science (JSPS KAKENHI) Grant Number 17H04195(MK, AH, YI)

JSPS KAKENHI Grant Number 20H00527(MK, YI)

JSPS KAKENHI Grant Number 21K15677(DI)

JSPS KAKENHI Grant Number 23K14773(DI)

JSPS KAKENHI Grant Number 23H00420(MK, MI, YI)

Japan Agency for Medical Research and Development (AMED) Grant Number JP21wm0425008(TY)

AMED Grant Number JP21wm0425013(MK)

AMED Grant Number JP22nk0101575(MK)

AMED Grant Number JP22am0401007(MK)

Research grant from Mitsubishi-Tanabe Pharma (MK)

## Author contributions

Daisuke Ito: Conceptualization, methodology, clinical investigation, molecular experiments, writing – original draft. Madoka Iida: Conceptualization, methodology, molecular experiments, single nucleus RNA-Seq, supervision, writing – review & editing. Yohei Iguchi: Conceptualization, methodology, supervision, writing – review & editing. Atsushi Hashizume: Methodology, clinical investigation, supervision. Shinichiro Yamada: Clinical investigation. Yoshiyuki Kishimoto: Clinical investigation. Shota Komori: Clinical investigation.Teppei Shimamura: Scientific discussions. Yuto Takemoto: Scientific discussions. Masahiro Nakatochi: Scientific discussions. Tomohiro Akashi: Single nucleus RNA-Seq. Kunihiko Hinohara: Single nucleus RNA-Seq. Hyeon-Cheol Lee-Okada: Lipidomics Junichi Niwa: Scientific discussions Gen Sobue: Scientific discussions. Shinji Tanaka: Scientific discussions. Ken Takashina: Scientific discussions. Takehiko Yokomizo: Lipidomics. Masahisa Katsuno: Conceptualization, methodology, supervision, writing – review & editing.

## The Paper Explained Problem

There is a considerable variability in the disease progression of sporadic ALS, but the molecular basis for phenotypic heterogeneity remains largely unknown. Herein, to elucidate the systemic metabolic changes related to ALS progression, we performed metabolomics analysis on the serum of ALS patients and identified several metabolites associated with the disease progression。

## Result

In this study, we clinically investigated metabolic changes related to the progression speed of ALS using metabolomics analysis of serum derived from ALS patients, revealing several metabolic dysregulation related to ALS progression, including N-acyl taurines which belong to the expanded endocannabinoid system. The administration of PF-04457845, a fatty acid amide hydrolase (FAAH) inhibitor which increases the expanded endocannabinoid system, rescued ameliorated the phenotype of the cellular models and a murine model of ALS.

## Impact

This study demonstrated that dysregulation of the expanded endocannabinoid system (ECS) contributes to the rapid progression of ALS and the ECS can be a promising target for ALS cure.

